# The causal direction in the association between Parkinson’s disease and cigarette or nicotine use

**DOI:** 10.1101/161240

**Authors:** Nikodem Grzesiak

## Abstract

The association between cigarette use and Parkinson’s disease (PD) has been well known in the literature since more than 50 years ago. Large population studies showed that smokers developed PD less often than non-smokers and that the duration of smoking was inversely proportional to PD risk. There are two primary hypotheses for this association in the literature. First, long-standing hypothesis, is that smoking cigarettes is neuroprotective. The second, recent hypothesis, is that there exists biological predisposition to PD, which also manifests in decreased stimulus seeking behavior, hence lesser likelihood to smoke or use nicotine (reverse causation). The objective of this article is to summarize the evidence available in the literature and evaluate the causality of the association between Parkinson’s disease and smoking cigarettes or nicotine ingestion. It is found that the first, directly causal hypothesis is a stronger contributor to the effect, although the reversely causal mechanism could play a role. It is found that smokeless tobacco use decreases the risk of PD stronger than smoking cigarettes does, suggesting that nicotine is more important in neuroprotection than other cigarette smoke constituents.

## I. Smoking cigarettes protects against Parkinson’s disease

The earliest association of the decreased risk of PD in smokers was discovered in 1959, in a collaborative study by the United States Public Health Service and the Veterans Administration. The study investigated the various causes of death of approximately 200, 000 U.S. government life insurance policyholders, who reported their tobacco use status and frequency. Smokers showed a dramatically lower mortality from PD. For those who smoked 10-20 cigarettes daily, the ratio of observed to expected deaths was 0.11 [Dorn, 1959.]. Since then many studies and meta-analyses have been conducted on the subject. A meta-analysis of four cohort studies and 44 case-control studies showed that compared to never smokers, the relative risk of PD was 0.80 [95% CI, 0.69-0.93] for former smokers (*>* 100 cigarettes in a lifetime), 0.59 [0.54-0.63] for ever smokers (*>* 100 cigarettes in a lifetime; still smoke, but not every day), and 0.39 [0.32-0.47] for current smokers [Hernan et al., 2002].

The analysis of longitudinal data from 305, 468 participants of the NIH-AARP Diet and Health cohort, of whom 1662 had a PD diagnosis, clearly showed a positive association between the number of years of smoking and the reduced risk of PD. Compared to never-smokers, the odds ratio (OR) was found to be 0.78 for past smokers and 0.56 for current smokers. Additionally, the longer duration of smoking corresponded to diminished OR of Parkinson’s. Regardless of the number of cigarettes smoked per day, the OR for those who smoked, for 1-9, 10-19, 20-29, 30-39, and 40+ years were respectively [95% CI]: 0.92 [0.79-1.07], 0.75 [0.63-0.88], 0.72 [0.61-0.85], 0.65 [0.53-0.79], 0.54 [0.41-0.70] [Chen et al., 2010].

**Figure 1:**
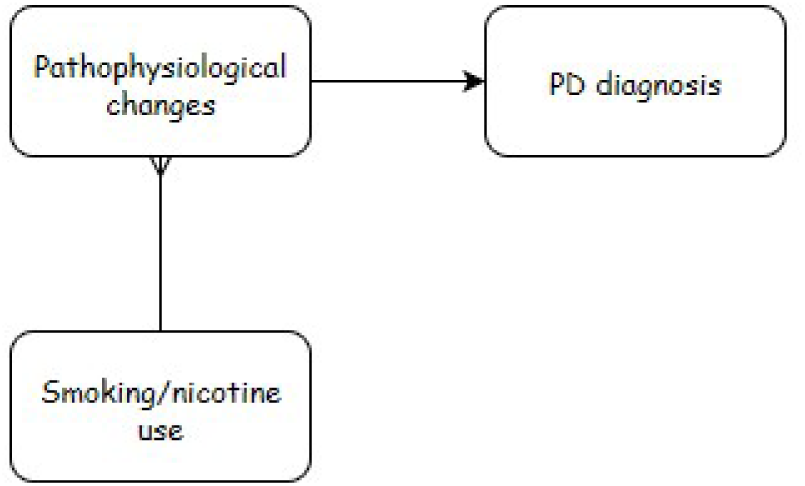
A potential causal model for smoking as a neuroprotective agent. The stimulatory arrow represents synergy, while the inhibitory arrow indicates antagonism.

These and numerous other studies put forward the following hypothesis: smoking is neuroprotective against PD; it slows down the development of PD pathophysiological changes.

## II. Parkinsonian predisposition alters the smoking behavior

Several authors of smaller studies investigating the association, interpreted the findings differently, observing that subjects with biological predisposition to PD display specific personality traits that, among others, make them less likely to smoke. In one study, for example, PD subjects scored 3.0, [95% CI, 0-8] compared to controls 3.9, [0-12] on the Sensation Seeking Scale. The authors hypothesized that these decreased risk taking and stimulus-seeking behaviors, as in, perhaps, coffee and alcohol consumption in PD patients are probably due to low levels of dopamine in the brain [Evans et al., 2006]. Hence, supposedly, the progressive depletion of dopaminergic neurons characteristic to PD, results in loss of nicotine “reward” or “sensitivity” that facilitates smoking cessation. Similarly, it has been suggested that loss or downregulation of nicotinic receptors precedes neurodegeneration [Perry et al., 1995].

A recent case-control study investigated the association of the ability to quit smoking and Parkinson’s disease in 1808 PD subjects and 1876 (sex and year of birth matched) controls, extracted from Danish records for years 1996-2009. The researchers used questionnaires to assess the difficulty of smoking cessation. They gathered qualitative data on the ability to quit smoking in four categories: “Easy”, “Moderate”, “Difficult but successful first attempt” (DBSFT) and “Extremely difficult”. They also asked about the use of nicotine substitutes (ever, never). Unsurprisingly, they found that smokers had a lower adjusted OR of PD, 0.65 [95% CI, 0.56-0.76] for former, and 0.28 [0.56-0.76] for current smokers. However, compared to those who quit “easily”, former smokers who found quitting “extremely difficult” had a risk of 0.69 [0.48-0.99], where the risk for both the DBSFT and moderate groups was 0.86 [0.64-1.14 and 0.63-1.19]. Having a hard time quitting smoking meant a 31% decreased risk of developing PD and over 9 times higher likelihood of using nicotine substitutes compared to those who quit “easily”. Authors suggested that because the subjects who developed PD had an easier time quitting smoking (and used less tobacco substitutes) than the controls, they were biologically predisposed to PD. Supposedly then, ease of quitting smoking is an early marker of PD, and the epidemiological relationship at hand is not a direct effect of smoking, but rather, a result of reverse causation [Ritz et al., 2014]. The reverse causation hypothesis is illustrated in Figure 2. The loss of sensitivity to stimuli decreases the odds of smoking and increases the risk of Parkinson’s disease.

**Figure 2:**
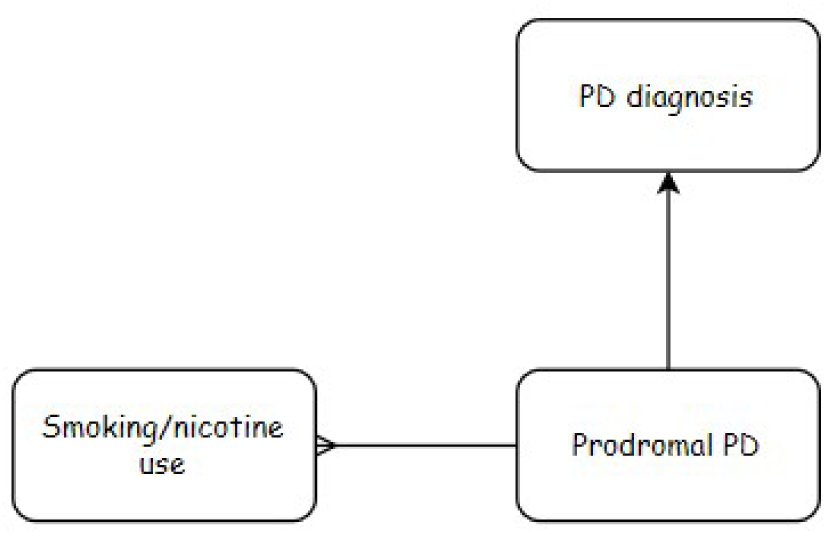
A potential model for prodromal PD (biological predisposition) affecting smoking / nicotine behavior

The authors hypothesized that the association is caused by genetic factors. Indeed, it has been documented that even though environmental factors like peer influences shape nicotine, alcohol, and caffeine use behaviors in adolescence, genetic factors become progressively more dominant later in life [Kendler et al., 2008]. Specific genes responsible for acetylcholine receptor subunits and nicotine metabolism have been found to modify nicotine sensitivity [Thorgeirsson et al., 2010]. Hence, a genetic link between nicotine sensitivity and PD is plausible, such that smoking behavior could predict PD risk.

Albeit an interesting observation, a more recent case-control study from 2014 aimed to quantify the protective effects of stimulants such as cigarettes and alcohol and caffeine consumption on Parkinson’s disease risk, using a novel modeling method. Unknowingly, the researchers found strong evidence supporting only one of the hypotheses. The data collected from 444 PD patients and 876 controls revealed that even though the PD cases were much less likely than controls to ever have smoked cigarettes (53.4% vs 72.3%), the ever regular coffee consumption (96.2% vs 97.7%) and regular alcohol consumption (76.6% vs 77.5%) wasn’t much different [van der Mark et al., 2014]. This finding suggests that the stimulation seeking isn’t significantly reduced in PD subjects. Additionally, the researchers analyzed the former smokers according to pack-years and time since cessation. In a graph showing how OR changes with Time-Since-Cessation plotted for 15 pack-years of total smoking, the OR increased almost linearly with the time passed since cessation. Current 15 pack-years smokers had an OR of 0.2 and the risk went up by about 0.2 for every 20 years since cessation. The trend was present in all the pack-years groups, with slower risk increase after cessation for smokers with more pack-years. By the model, smoking a pack a day for 8.5 years decreases the OR of Parkinson’s disease by 80%. Also, 21 years after cessation it makes little difference if one smoked 20 Pack-Years or 100 Pack-Years (5% OR difference) [van der Mark et al., 2014]. This finding suggests that the duration of smoking and time-since-cessation are more relevant to PD risk than smoking intensity. This is also criticism to almost all the previous studies, where these crucial group distinctions were lost under a common “former smoker” grouping. Even though the prodromal (biological predisposition to) PD may play a role in the relationship at hand, there is a clear direct or indirect causal link between smoking and the disease onset. Once smoking is stopped, the OR progressively increases, which means that regardless of biological predisposition, smoking directly or indirectly affects the likelihood of the disease. The trend of PD odds ratio increase following smoking cessation suggests that smoking is neuroprotective.

**Figure 3:**
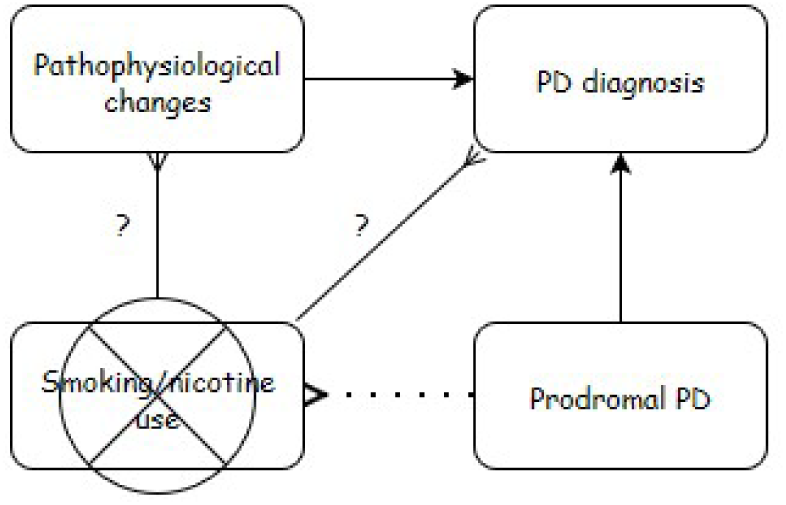
When active smoking is ceased, the OR of PD progressively increases. This suggests a limited influence of the prodromal PD. Smoking affects PD risk, through a simple or complicated mechanism (marked “?”).

## III. Other Evidence-Environmental Tobacco Smoke (ETS)

Other publications support the hypothesis that smoking is neuroprotective against Parkinson’s disease. Especially three ETS studies, one twin study and one population study seem to point towards the causality of the relationship. The ETS relationship to PD was studied as a proxy, since the effect of second-hand exposure to tobacco smoke (passive smoking) is independent of active smoking and therefore circumvents the sensation-seeking hypothesis. Regardless if the group were smokers or non-smokers, a measurable association between ETS and Parkinson’s disease would be evidence for a direct connection between them, as opposed to the reversely causal link, as in Figure 2.

### i. ETS at the workplace and home

The first case-control study of association of ETS and PD was conducted in 2006 on 163 Australian PD patients. The participants were asked about their smoking history with information about initiation, cessation, frequency and quantities smoked was collected. Regular smoking was defined as “smoking as often as once per week for 6 months or more”. Next both smokers and non-smokers were asked about the prevalence of ETS smoke in their life-either at the workplace or home. The data for both never and ever smokers who lived with a smoker or worked in a smoky workplace showed decreased ORs of PD. For example, the never smokers who lived with a smoker for 1-15, 15-25, 26+ years had the adjusted ORs [95% CI] of 0.67 [0.15-2.97], 0.71 [0.18-2.81] and 0.51 [0.07-3.67], respectively, compared to those who never lived with a smoker. The ETS exposure effect was dose-dependent-longer exposure further reduced risk for PD. [Mellick et al., 2006].

The 2012 ETS case-control study from the U.S. investigated the association between PD and living or working with active smokers in 153 PD subjects. Authors observed that compared to controls, passive never-smokers (exposed either at home or work) had a pooled OR of 0.34 [95% CI: 0.16-0.73] whereas the active smokers showed a similar OR of 0.35 [0.17-0.73] [Searles Nielsen et al., 2012]. The risk was again inversely proportional to years of ETS exposure.

Another case-control study on 249 PD patients carried out in Japan showed similar relationships for smokers (adjusted OR as low as 0.12 for current smokers), but no significant association between ETS exposure and PD (OR: 0.99) [Tanaka et al., 2010].

### ii. Parental smoking

To further explore the relationship between smoking and PD, researchers at Harvard, explored how parental smoking during childhood influenced PD incidence in the Nurses’ Health Study (26 years long) and the Health Professionals Follow-up Study (18 years long) in the U.S. During their respective durations, 455 new PD cases provided parental smoking information for the analysis. Compared to those whose parents didn’t smoke, the combined, age-adjusted, relative risk of PD was 0.73 (95% CI, 0.53, 1.00) for participants who reported that both parents smoked and 0.87 (95% CI, 0.64, 1.20) when one parent smoked. The caffeine and alcohol adjustments didn’t significantly alter the results. [O’Reilly et al., 2009].

The results from the passive smoking studies, support the hypothesis that ETS, independent of active smoking, reduces the risk of Parkinson’s disease.

**Figure 4:**
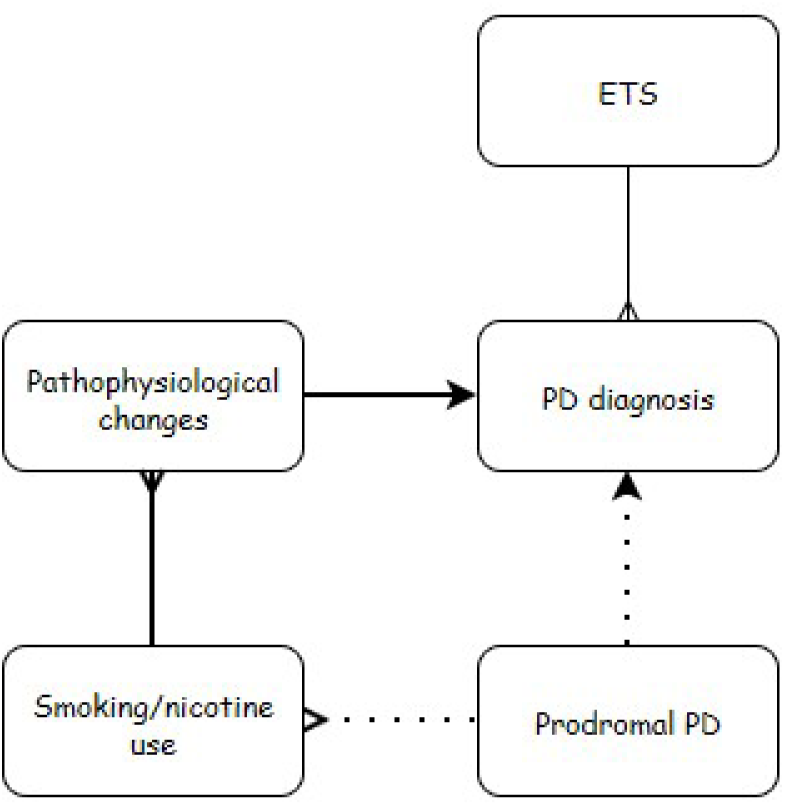
A model of smoking / PD relationship. ETS exposure as an instrumental variable for testing the causality, suggests the neuroprotective effect of smoking / nicotine. The dotted link represents weak association and the bolded link represents strong association.

### iii. Twin studies

According to the inversely causal hypothesis, the relationship between smoking and PD could be the result of a genetic factor related to both smoking behavior and the chance of developing PD. The twin studies, which circumvent the influence of genetic and, to a lesser degree, environmental cofounding in the relationship, showed that this isn’t the case. Even though the total sample size was relatively small (528 twin pairs in 2 studies), the smoking twins had a lower incidence of PD than their nonsmoking counterparts [Tanner et al., 2002];[Wirdefeldt et al., 2005]. Among twin pairs, the twin who developed PD stopped smoking on average 3.7 years earlier than the twin without PD; when both developed PD, the one affected first stopped smoking on average 1.3 years earlier [Tanner et al., 2002].

### iv. Population trends

Data since 1860 shows, that the percentage of ever-smoking women has been increasing, while the same statistic for men has been on a decline in both the UK and the U.S. Using relevant literature, the researchers estimated the male to female ratios in PD incidence and correlated them with the corresponding male to female ever-smoking prevalence ratios, obtained from national records. In result, they showed that the relative frequency of PD among women declined when the proportion of women smoking increased. The strength of this correlation (*r* = 0.28; *P* = 0.0002) is comparable to the opposite trend observed for lung cancer incidence in smokers. By the correlation smokers experienced a 74% PD risk decrease [Morozova et al., 2008].

A more recent study examined the trends in the incidence of parkinsonism and PD over 30 years in a geographically defined American population of 906 cases. Researchers found that the overall incidence rates increased significantly during that period, especially for men over 70. The IR increased by 1.24 per decade [1.07-1.44] for parkinsonism and by 1.35 per decade [1.10-1.65] for PD. In conclusion the authors suggested that these trends may be associated with the dramatic changes in smoking behavior or other environmental factors that took place in the second half of the 20th century [Savica et al., 2016].

### v. Imaging studies

There is another line of evidence opposing the stimuli aversion hypothesis in prodromal PD subjects. Imaging studies have shown nigrostriatal dopaminergic neuronal loss starting to increase less than 10 years before onset of clinical symptoms of PD, suggesting a limited impact of reduced sensation seeking earlier in life [Savica et al., 2010].

### vi. Other possible causation models

The association could be explained by a still-unknown third factor that increases the risk of Parkinson’s disease and causes an aversion to cigarettes or nicotine (U in the figure below).

**Figure 5:**
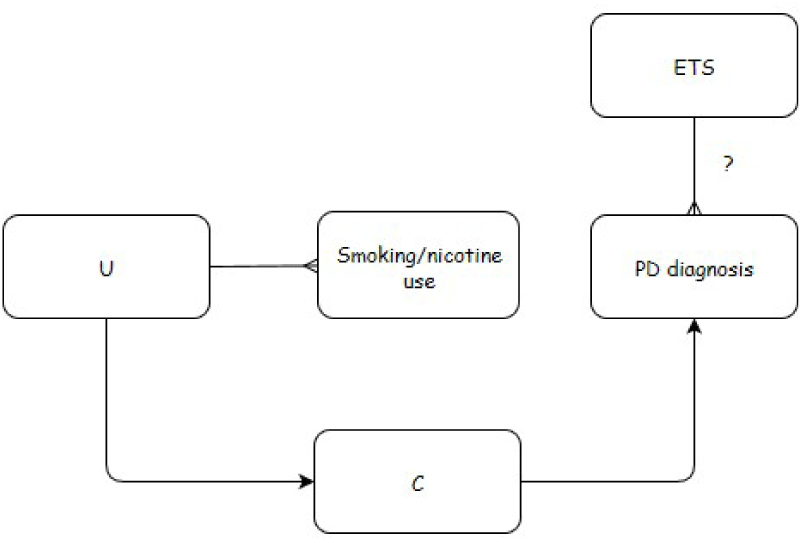
Another possible causal model for the relationship. U is an independent predictor (i.e. some toxin) that inhibits smoking and links to some confounding factor that causes PD. Findings from ETS studies are hard to explain under this hypothesis.

This factor is unlikely to be genetic, because the twin studies showed a direct connection between smoking and PD risk. However, confounding by an environmental factor cannot be eliminated by twin studies. For example, a person could be exposed to a toxic chemical or an infectious agent that causes subclinical damage to the dopaminergic system, which is involved in novelty seeking and addiction, and this exposure could reduce their desire to smoke while independently increasing the risk of Parkinson’s disease. However, if the association between smoking and Parkinson’s disease were solely due to exposure to a toxic chemical or an infectious agent, no relation would be expected between ETS and Parkinson’s disease [Searles Nielsen et al., 2012]. Additionally, the strong association of population-wide smoking rates and PD incidence make this causation model unlikely.

## IV. Nicotine as most important in the association

Tobacco smoke contains thousands of substances any one of which could be responsible for the evident association between PD and smoking. Nicotine is usually regarded as the most important one for a few reasons. Nicotine acts at the nicotinic acetylcholine receptors which have a close anatomical relationship with the dopaminergic neurotransmitter system in the striatum whose neurons are selectively damaged in PD. Also, nicotine has been found protective against neuronal damage in cellular and animal models of PD [Parain et al., 2003]. To test the association between nicotine and PD, we can explore a less contaminated source of the compound available-smokeless tobacco (ST). ST products contain many fewer substances and provide the users with similar amount of nicotine as smoking cigarettes does, roughly 1mg [Benowitz et al., 1988]. There are three ST studies and one dietary nicotine study relevant to the nicotine-PD association. One case-control study on 196 PD subjects found significantly reduced OR [95% CI] of 0.18 [0.04-0.82] in ever users vs. never users of ST [Benedetti et al., 2000]. Another, prospective cohort study assessed PD mortality as the outcome (by death certificate) in 95, 981 never-smoking men and found a relative multivariate adjusted risk of 0.24 [0.08-0.75] for current users of smokeless tobacco vs. never users [O’Reilly et al., 2005]. The strongest evidence comes from a recent study with a sample of 348, 601 men. Researchers combined data from seven prospective cohort studies and used survival analysis with multivariable Cox regression to find the relative change in PD risk due to ST use, restricting their analysis to never-smokers to limit confounding. They found that compared with never-ST users, the ever-ST users had a pooled risk of 0.41 [95% CI 0.28-0.61] [Yang et al., 2016].

Interestingly, foods from the same botanical family as tobacco, Solanaceae, contain small amounts of edible nicotine: on average 102.1, 43.8 and 19.25 *µ*g/kg for pepper, tomato and potato, respectively. In a case-control study with 486 PD cases, researchers collected data on the tobacco use histories and frequency of consumption of Solanaceae family edibles and 17 other vegetables as for comparison. They found that the consumption of dietary nicotine was inversely related to PD risk, and the association strengthened when weighted by nicotine concentration. There was an inverse association for peppers with OR of 0.59 [95% CI: 0.35-0.99] for never-users of tobacco, who consumed them 2-4 times / week [Nielsen et al., 2013]. Like ETS and others, this study adds support to the neuroprotective as opposed to the reverse causation explanation of smoking / nicotine-PD association.

The four studies are evidence that nicotine is an important neuroprotective constituent of cigarette smoke. Furthermore, since the protection from ST, a less contaminated nicotine source of nicotine, seems to be stronger than that due to smoking, perhaps some of the toxic substances in the smoke diminish the positive neuroprotective effects of nicotine.

## V. Possible mechanism of action-Low concentration hypothesis

In light of the ETS and food studies, some researchers studying the causality suggested that the duration of exposure (even at low concentrations) might be the most important neuroprotective factor against PD. Honglei Chen, a senior researcher at the National Institute of Environmental Health Sciences and a co-author of many smoking-PD studies, wrote that it is possible that the biological effects of nicotine or other tobacco compounds may be saturated at low concentrations. He added that since PD likely takes time to develop, sustained exposure may be needed to lower the risk [Chen, 2015]. Other researchers proposed that smoking might have a protective effect not through direct action of a specific compound, but rather because chronic low dose exposure to smoke toxins can increase tolerance to acute exposures of other toxic compounds like *α*-synuclein and A*β*, which cause PD neurodegeneration [Yang et al., 2016].

## VI. Conclusion

Having examined the evidence, we can conclude strong support to the causal neuroprotective relationship of smoking to Parkinson’s disease. The pathophysiological changes and the prodromal PD boxes may very well have many of the same underlying factors, like similar mediation mechanisms and brain areas involved. The insensitivity to nicotine may have some backwards effect on the smoking behavior, but the neuroprotective action of nicotine certainly seems dominant in the association.

**Figure 6:**
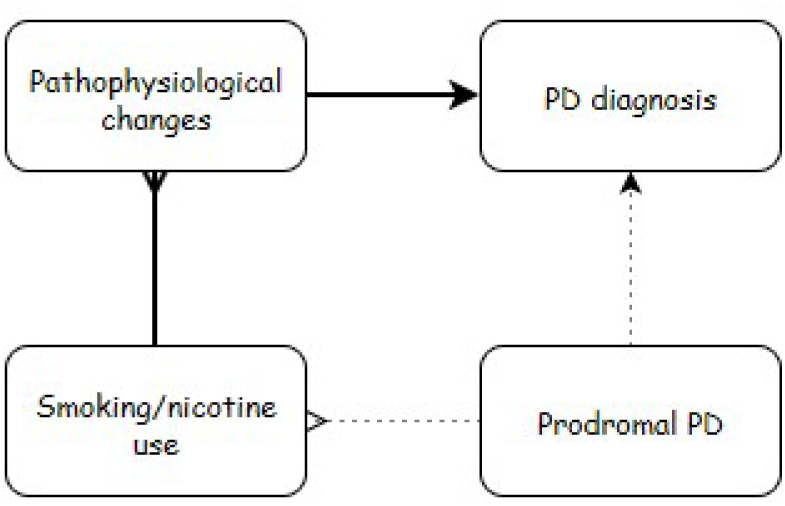
Evidence shows that the causal (neuroprotective) association contributes a greater effect that the reversed causal association.

